# Fractionation of impulsive and compulsive trans-diagnostic phenotypes and their longitudinal associations

**DOI:** 10.1101/570218

**Authors:** Samuel R. Chamberlain, Jeggan Tiego, Leonardo F. Fontenelle, Roxanne Hook, Linden Parkes, Rebecca Segrave, Tobias U. Hauser, Ray J. Dolan, Ian M. Goodyer, Ed Bullmore, Jon E. Grant, Murat Yücel

**Affiliations:** Department of Psychiatry, University of Cambridge, and Cambridge and Peterborough NHS Foundation Trust, UK; Monash Institute of Cognitive and Clinical Neurosciences, School of Psychological Sciences, Monash University, c/o MBI, 770 Blackburn Rd, Clayton, VIC. 3800, Australia; The Max Planck UCL Centre for Computational Psychiatry and Ageing, University College London (UCL), UK; Department of Psychiatry & Behavioural Neuroscience, University of Chicago, USA

**Author notes:** Address correspondence (including reprint requests) to: Samuel R. Chamberlain, MD, PhD, MRCPsych, Department of Psychiatry, University of Cambridge, Addenbrooke’s Hospital, Cambridge. CB2 0QQ. United Kingdom.

## Abstract

**Objective:** Young adulthood is a crucial neurodevelopmental period during which impulsive and compulsive problem behaviours commonly emerge. While traditionally considered diametrically opposed, impulsive and compulsive symptoms tend to co-occur. The objectives of this study were: (i) to identify the optimal trans-diagnostic structural framework for measuring impulsive and compulsive problem behaviours; and (ii) to use this optimal framework to identify common/distinct antecedents of these latent phenotypes.

**Methods:** 654 young adults were recruited as part of the Neuroscience in Psychiatry Network (NSPN), a population-based cohort in the United Kingdom. The optimal trans-diagnostic structural model capturing 33 types of impulsive and compulsive problem behaviours was identified. Baseline predictors of subsequent impulsive and compulsive trans-diagnostic phenotypes were characterised, along with cross-sectional associations, using Partial Least Squares (PLS).

**Results:** Current problem behaviours were optimally explained by a bi-factor model, which yielded dissociable measures of impulsivity and compulsivity, as well as a general disinhibition factor. Impulsive problem behaviours were significantly explained by prior antisocial and impulsive personality traits, male gender, general distress, perceived dysfunctional parenting, and teasing/arguments within friendships. Compulsive problem behaviours were significantly explained by prior compulsive traits, and female gender.

**Conclusions:** This study demonstrates that trans-diagnostic phenotypes of 33 impulsive and compulsive problem behaviours are identifiable in young adults, utilizing a bi-factor model based on responses to a single questionnaire. Furthermore, these phenotypes have different antecedents. The findings yield a new framework for fractionating impulsivity and compulsivity; and suggest different early intervention targets to avert emergence of problem behaviours. This framework may be useful for future biological and clinical dissection of impulsivity and compulsivity.

## Introduction

Young adults are particularly vulnerable to develop maladaptive, problem, behaviours (Spear, 2000). This vulnerability is thought to stem from neurodevelopmental alterations (Casey et al., 2017) coupled with concomitant changes in the environment (such as lessening parental oversight and increasing exposure to new peer groups). Two key concepts of particular relevance to understanding brain development and problem behaviours in young adulthood are ‘impulsivity’ and ‘compulsivity’ (Hollander et al., 2009). Impulsivity refers to behaviours that are unduly hasty, risky, and that lead to negative outcomes in the long term (Evenden, 1999). Examples of clinical disorders characterised by impulsivity include antisocial personality disorder, gambling disorder, and substance use disorders, e.g. (Krmpotich et al., 2015). Thus an individual may undertake a spontaneous aggressive act (as in antisocial personality disorder), or may crave and consume a substance (or engage in a gambling opportunity), without due regard to potentially damaging consequences. Compulsivity, by contrast, refers to functionally impairing behaviours undertaken repeatedly, often according to rigid rules, manifesting classically in obsessive-compulsive disorder (OCD). Impulsive and compulsive problems are common in young adulthood, have negative effects on long-term outcomes, and frequently co-occur (Fineberg et al., 2013; Cerda et al., 2016; Black et al., 2017).

Mechanistic, diagnostic, and treatment progress in psychiatry has been hindered by an excessive focus on specific mental disorders, typically examined in isolation within clinical settings rather than a continuous or dimensional fashion in the population at large (Cuthbert and Insel, 2013). Intermediate phenotypes (trans-diagnostic markers) hold promise in this regard, and should be measurable along a continuum in the background population, existing in more extreme forms in people with conceptually related mental disorders. Recent reports from population-based studies have suggested that trans-diagnostic phenotypes are indeed likely (St Clair et al., 2017; Stochl et al., 2015). Thus, a crucial next step in impulsivity-compulsivity research and practice is to identify intermediate phenotypes that cut across related problem behaviours, and that are measurable dimensionally both in the general population and in the context of existing categories of mental disorders.

Recent cross-sectional studies have shown that variation in a broad range of impulsive and compulsive behaviours is also present sub-clinically, and that these behaviours can be grouped into latent dimensions of ‘impulsivity’ and ‘compulsivity’ (termed intermediate phenotypes), which are found to be positively correlated (Chamberlain et al., 2017; Guo et al., 2017). This positive correlation could reflect common underpinnings; but also, could theoretically stem from common measurement bias (cross-talk across scale items due to item response bias). It is important to identify an optimal framework within which to conceptualize separable impulsive and compulsive problem behaviours as this could then be used to explore common or distinct antecedents. A recent resurgence of interest in bi-factor models in psychiatry (Reise, 2012b; St Clair et al., 2017), has yet to be adequately deployed and tested in latent phenotyping studies of impulsivity and compulsivity (Chamberlain et al., 2017; Guo et al., 2017). Such bi-factor models incorporate a general factor (capturing common variance across all study measures), as well as specific factors, enabling – theoretically – these constructs to be truly fractionated statistically.

The aims of the current study, were: (i) to identify the optimal trans-diagnostic structural framework for measuring impulsive and compulsive problem behaviours (including consideration of a bi-factor model); and (ii) to use this optimal framework to identify longitudinal antecedents of these latent phenotypes for the first time, with a particular focus on demographic, personality, parenting, and friendship measures. We focused on these measures as they were expected to contribute to subsequent problems, based on prior evidence. The optimal conceptual framework was identified using a competing models approach; and relationships between trans-diagnostic phenotypes and other measures were elicited using the innovative statistical approach of Partial Least Squares, which is well-suited for data likely to be correlated and non-normally distributed (Abdi and Williams, 2013).

## Method

### Participants

We contacted all individuals from the Neuroscience in Psychiatry Network (NSPN), a longitudinal population-based cohort study examining brain development (Kiddle et al., 2017), via email. Participants completed an online survey implemented in SurveyMonkey in 2018 and had also provided baseline questionnaires via the post (2012-2014). The current data collection (2018) examined a broader range of impulsivity/compulsivity measures, some of which were not available at the time the NSPN cohort was conceived (Chamberlain and Grant, 2018; Guo et al., 2017). The study was approved by Research Ethics Committee and individuals provided informed consent. Participants were given a £15 voucher for taking part.

### Measures of interest

Questionnaires collected presently (2018) were designed to capture relevant demographic characteristics (age, gender, ethnic group, level of education), trans-diagnostic measures of problem behaviours, and other relevant measures of impulsivity and compulsivity; plus overall quality of life. The outcome instrument of interest, used to examine underlying latent phenotypes of impulsive and compulsive problem behaviours, was the Impulsive-Compulsive Behaviours Checklist (ICBC) (Guo et al., 2017), which is described in more detail below. Other measures of interest collected in the current data round (2018) and at participant recruitment (2012-2014) are described in **Table 1**. The latter included measures of impulsive-compulsive tendencies, but also parenting, and friendships, which we expected may impact the ultimate expression of the latent phenotypes on the ICBC. Additionally, the following demographic information was examined, which was completed by the participant’s parent or legal guardian at study baseline: any perinatal complications in proband (yes/no), any history of head trauma in proband (yes/no), family educational level (parent/guardian’s number of years’ education completed), and any current or previous emotional/behavioural/mental health problem in the proband (yes/no).

The ICBC enquires about the presence of 33 types of impulsive and compulsive problem behaviours; for each type of behaviour, the individual endorses whether they and/or others think they have a problem with the behaviour, responding: never, sometimes, often, or always. The behaviours asked about by the ICBC are as follows: washing, smoking, feeling compelled to collect things, being overly cautious with money, re-arranging/ordering, shopping, list making, counting (e.g. money, tiles), grooming, idiosyncratic routines (performing a very personalised sequence of actions), repeating actions (over and over again), exercising, betting/gambling, hair pulling, lying, sexual activities/behaviours, alcohol consumption, planning (e.g. over-organising), illicit drug use, cleaning too much, verbal aggression, violence towards objects/properties, swearing, checking (e.g. locks, light switches), checking (e.g. yourself in the mirror), speed driving, medication use, physical aggression, social networking (e.g. Facebook, twitter, Google+, Myspace), applying rules, purposeful self-injury (i.e. not accidental), re-writing/re-reading, and tattooing. Previous work found the scale to have sound psychometric properties (Guo et al., 2017).

**Table 1.** Summary of instruments included in the study. Full details for the instrument citations are provided in the online supplement, due to space limitations.

| **Instrument** | **Description** | **Outcome measure(s)** |
| --- | --- | --- |
| *Measures collected in current data round* |  |  |
| Barratt Impulsiveness Scale (BIS-11 Brief version) (Steinberg, Sharp et al. 2013, Stanford, Mathias et al. 2016) | Barratt Impulsiveness Scale (BIS-11 Brief version). The BIS is an extensively used scale used for the classic measurement of impulsivity. We used the 8-item version, which has been previously validated in several studies. The questionnaire asks about impulsive tendencies and behaviours; for each item, the individual indicates the extent to which it applies to them (not at all, a little, quite a lot, or very much; scored 0-4 per item) | Total score |
| Antisocial personality disorder symptoms (American Psychiatric Association 2013) | Antisocial personality disorder symptoms were quantified using a tick-list of DSM-5 criteria. For each symptom criterion, participants indicated whether the item applied to them or not | Total number of criteria endorsed |
| Padua obsessive-compulsive inventory (Washington State University Revision version) (Sanavio 1988, Burns 1995) | This is a 39-item questionnaire measuring obsessive-compulsive symptoms in normative and clinical settings | Symptom domain scores (per Burns et al.) |
| Cambridge-Chicago Trait Compulsivity Scale (CHIT) (Chamberlain and Grant 2018) | This is a 15-item scale capturing day-to-day aspects of compulsivity. For each item, the individual indicates whether the statement (e.g. "I hate leaving a task unfinished") applies to them. Response options are: strongly disagree (0), disagree (1), agree (2), or strongly agree (3) | Total score |
| Obsessive Beliefs Questionnaire Short Form | This is a 20-item short version of a widely used questionnaire to assess obsessional beliefs in the context of OCD. For each item, the person indicates how much they | Summary scores: threat perception, perfection/intolerance of uncertainty, |
| (OBQ) (Moulding, Anglim et al. 2011) | agree with a given statement, ranging from disagree very much (scored 1) to agree very much (7). | inflated responsibility, and importance/control of thoughts |
| Obsessive-Compulsive Personality Disorder traits (OCPD) (Sheehan, Lecrubier et al. 1998, American Psychiatric Association 2013) | The diagnostic screening questions from the Mini-International Neuropsychiatric Inventory (MINI) were used for OCPD traits. For each item, participants were asked whether the statement applied to them, taking into consideration not only their own view but also what they felt other people (close confidantes) would think | Total number of OCPD traits endorsed |
| General psychopathology (depression/anxiety) from the Mini-International Neuropsychiatric Inventory (MINI) (Sheehan, Lecrubier et al. 1998) | General psychopathology (depression/anxiety). The diagnostic screening questions from the Mini-International Neuropsychiatric Inventory (MINI) were used for depressive and anxiety disorders | Presence or depressive and/or anxiety symptoms (total items endorsed) |
| Brunnsviken Brief Quality of life scale (BBQ) (Lindner, Frykheden et al. 2016) | The BBQ is a previously validated self-report quality of life scale, which covers six life areas (leisure time, view of life, creativity, learning, friends/friendship, and view of self) that are important determinants of overall quality of life. This scale has been validated in various normative and clinical population settings | Total quality of life score |
| *Measures collected at study baseline (2012-2014)* |  |  |
| Barratt Impulsiveness Scale (BIS 11 full version) (Patton, Stanford et al. 1995) | This question asks about impulsive tendencies and behaviours; for each, the individual indicates the extent to which it applies to them (not at all, a little, quite a lot, or very much; scored 0-4 per item) | Total score |
| The Antisocial Process Screening Device (APSD) (Munoz and Frick 2007) | This is a 20-item questionnaire developed to assess psychopathy in young people. For each item, the individual indicates whether a given statement (e.g. "You engage in illegal activities") is not true at all (0), sometimes true (1), or definitely true (2) | Total score |
| Padua obsessive-compulsive inventory (Washington State University Revision) | Per current data round | Per current data round |
| version) (Sanavio 1988, Burns 1995) |  |  |
| General psychopathology (K10) (Kessler, Andrews et al. 2002) | The K10 was developed for assessment of generalised psychological distress, including distress arising from anxiety and depression symptoms. There are ten items each asking about experiences over the preceding 30 days. For each item, responses are: none of the time (0), a little of the time (1), some of the time (2), most of the time (3), or all of the time (4) | Total score |
| Family Assessment Device (FAD) (Ridenour, Daley et al. 1999), Positive Parenting Questionnaire (PPQ), Alabama Parenting questionnaire Child Short Form (APQ) (Elgar, Waschbusch et al. 2007) and Measure of Parenting style (MOPs) (Parker, Roussos et al. 1997) | Several complementary questionnaires relating to parenting approaches were included. The reader is referred to the cited manuscripts for more detailed descriptions of these instruments, except for the PPQ, which was developed bespoke for the NPSN study. The PPQ asks the extent to which each of 26 positively worded items related to one's family when living at home (e.g. "We spent quality time together."; "If I was angry, I was still listened to.") Each item is responded to with always (scored 0), mostly (1), sometimes (2), or rarely (3). Exploratory Factor Analysis (EFA) was used to identify the optimal data structure across all these questionnaires' items considered collectively (**Supplementary Table 2**) | Factor scores: general parenting, dysfunctional parenting (paternal side), and dysfunctional parenting (maternal side) |
| Friendship questionnaire (Goodyer, Herbert et al. 1997, van Harmelen, Kievit et al. 2017) | This questionnaire assesses the number, availability and adequacy of friendships as well as self-disclosure and difficulties in friendships, and was adapted from previous research work. The individual questions were: are you happy with the number of friends you have? How often do you arrange to see friends other than at school, college or work? Do you feel that your friends understand you? Can you confide in your friends? Do your friends ever laugh at you or tease you in a hurtful way? Do people who aren't your friends laugh at you or tease you in a hurtful way? Do you have arguments with your friends that upset you? Overall, how happy are you with your friendships? Item response options differed depending on the question but involved 4-6 possible responses (for example, for the question "Do you feel that your friends | Factor scores: happiness/number of friends, confiding/understanding, and teasing/arguments |
|  | understand you?”, the response options were most of the time, sometimes, not often, or not at all, which were scored 0, 1, 2, or 3 respectively). Exploratory Factor Analysis (EFA) was used to determine the latent structure of the questionnaire (**Supplementary Table 3**). |  |

(Note: references below are included in the supplement due to space limitations in the main manuscript)

### Data analysis

All data analyses were conducted using JMP Pro Version 13.2.0 (SAS Institute Inc.) and Mplus 7.2 (Muthén and Muthén, 1998 - 2012). The latent factor structure of the Impulsive-Compulsive Behaviours Checklist (ICBC, the outcome instrument of interest) was examined using Confirmatory Factor Analysis (CFA) of the covariance matrix using the Weighted Least Squares Means and Variance (WLSMV) adjusted estimator and Theta parameterisation (Muthén et al., 1997; Muthén and Muthén, 1998 - 2012). Based on prior studies and existing conceptual frameworks (Guo et al., 2017; Chamberlain et al., 2017; Reise, 2012a) we evaluated: a correlated two-factor model (two latent factors, i.e. impulsivity and compulsivity, correlated with each other), an orthogonal two-factor model (two latent factors, i.e. impulsivity and compulsivity, not correlated with each other), a one-factor model (one latent factor, corresponding to disinhibition), and a bi-factor model (two latent factors, i.e. impulsivity and compulsivity, plus a general factor). *Post hoc* model fitting was conducted by freeing theoretically plausible error covariances for estimation one at a time with reference to the highest modification index and these were corrected for significance using the Benjamini-Hochberg False Discovery Rate (.05). Test of exact model fit using the chi square test statistic (χ^2^) is overly sensitive to minor model misspecification in large sample sizes (Kline, 2016). Model fit was therefore evaluated using a combination of comparative fit indices, including the Root Mean Square Error of Approximation (RMSEA) (ε < .05 close approximate fit; ε = .05 - .08 reasonable approximate fit; ε = > 1.0 not close approximate fit); Comparative Fit Index (CFI) (> .90 reasonable fit; >.95 good fit); and the Weighted Root Mean Residual (WRMR) (<.950 good fit). We used a competing models approach, in which the comparative fit of several alternative models were evaluated to determine which provided the best representation of the covariances in the data.

Relationships between explanatory variables of interest and current impulsive-compulsive behaviours (latent scores on the ICBC) were investigated in two separate statistical models: the first examined baseline explanatory variables collected at study entry approximately (on average) 4 years prior; and the second model examined current explanatory variables (the broader range of impulsive and compulsive measures collected via the Internet in 2018). The statistical method of Partial Least Squares (PLS) was used. PLS is a versatile multivariate approach to data modelling that analyses relationships between exploratory (X) variables and outcome (Y) variables by means of fitting one or more latent components (Abdi and Williams, 2013). Unlike standard regression, PLS is robust to violations of normality assumptions and to item cross-correlations. Hence PLS is ideally suited to the current dataset. Candidate explanatory (X) variables in the PLS models were summary scores from the questionnaires, and the outcome variables of interest (Y) were the ICBC latent scores.

For the model exploring baseline predictors of later impulsive and compulsive problem behaviours, the X variables of interest were: age at baseline, gender, ethnic group, family education level (average years’ education for main caregiver), current or past mental health or behavioural problem (parent/guardian report), history of head trauma (parent/guardian report), history of any medical conditions (parent/guardian report), Barratt Impulsiveness Scale scores, antisocial personality total scores, Padua Inventory obsessive-compulsive scores, general psychopathology (K10 scores, including anxiety/depression), parenting scores (general parenting, dysfunctional paternal parenting, and dysfunctional maternal parenting), and friendship scores (happiness/number, confiding/understanding, and teasing/arguments). For the model exploring current measures potentially associated with problem behaviours, X variables of interest were: age, gender, education level, ethnic group, Barratt Impulsivity Scale scores, antisocial personality disorder traits (number of diagnostic criteria met), Padua Inventory obsessive-compulsive scores, OBQ scores, CHIT compulsivity scores, OCPD traits (number of OCPD criteria met), and general psychopathology (presence of depression or anxiety symptoms).

For PLS, missing data points were imputed automatically by JMP software using mean substitution. The PLS models were fitted using leave-one-out cross-validation (non-linear iterative partial least squares, NIPALS algorithm), and the optimal model was identified based on minimizing predictive residual sum of the squares (PRESS). From each initial model, measures with a Variable Importance Parameter (VIP) <0.8 were excluded per convention (Cox and Gaudard, 2013).

Explanatory (*X)* variables significantly contributing to the model (i.e., explaining significant variance in current compulsive and impulsive problem behaviours) were identified on the basis of 95% confidence intervals for bootstrap distribution of the standardised model coefficients not crossing zero (*N* = 1000 bootstraps; p<0.05).

## Results

Six-hundred and fifty-four individuals completed the study. The mean (standard deviation) current age was 23.4 (3.2) years, and 34.7% were males. The current Barratt Impulsiveness Scale (BIS-11 short) mean total score was 16.4 (3.9) and the Padua Inventory mean score was 19.0 (19.1). Participants had returned their baseline data to NSPN on average 4.0 (1.3) years prior to participation in the current study, and were of mean age 19.5 (3.1) years at baseline.

The majority of individuals (354 [53.2%]) endorsed having at least one problem behaviour in the preceding 12 months to some degree, on the ICBC (**Supplementary Table 1**). A bi-factor model consisting of a general latent ‘Disinhibition’ factor with loadings from all 33 ICBC items and two specific factors capturing residual variance in 14 impulsivity (“Impulsivity”) and 11 compulsivity items (“Compulsivity”) provided the best overall fit to the ICBC data compared to three competing models (**Table 2**). The loadings of individual behaviours onto the latent factors are shown in **Table 3**, and the model had had excellent fit (Reise, 2012b). Bi-factor models have a tendency to provide superior fit compared to correlated factors and indirect hierarchical models because they have more parameters (Bonifay et al., 2017). Model diagnostics, including measures of construct replicability, reliability, and unidimensionality were conducted to ensure the general and group factors were capturing meaningful covariances in the data and were not simply a product of overfitting (Rodriguez et al., 2016b). Construct replicability was evaluated by calculating the *H* index for the Disinhibition (.95), Impulsivity (.83), and Compulsivity (.69) factors, which ranges from 0 – 1 and quantifies the proportion of variance in a factor captured by its indicator variables (Hancock and Mueller, 2001). The results indicate that the strength of the standardised loadings for ICBC items on the latent factors were strong enough to suggest moderate-high replicability across studies using the same measures (Hancock and Mueller, 2001; Rodriguez et al., 2016b).

Calculation of the Explained Common Variance (ECV) revealed that the Impulsivity and Compulsivity group factors collectively accounted for sufficient explained variance (31%) in the ICBC, in addition to the general Disinhibition factor (69%), to justify retaining them over a unidimensional model (Rodriguez et al., 2016a; Reise, 2012b). Information functions generated for the 33 ICBC items loading on the general Disinhibition factor and were relatively distinct for different items (**Supplementary Figure 1**), suggesting that the general factor was not simply capturing item response bias or other forms of spurious variance (Bonifay et al., 2017). Factor score estimates were generated for the Disinhibition, Impulsivity, and Compulsivity factors for use in subsequent analyses, and exhibited weak bivariate correlations (Disinhibition with Impulsivity *r* = .115, *p* =.004 & Compulsivity *r* = .218, *p* <.001; Impulsivity with Compulsivity r = -.177, *p* < .001). These low correlations indicated that the factor score estimates for each latent variable were not heavily contaminated by variance from the other two factors (Grice, 2001).

**Figure 1.**
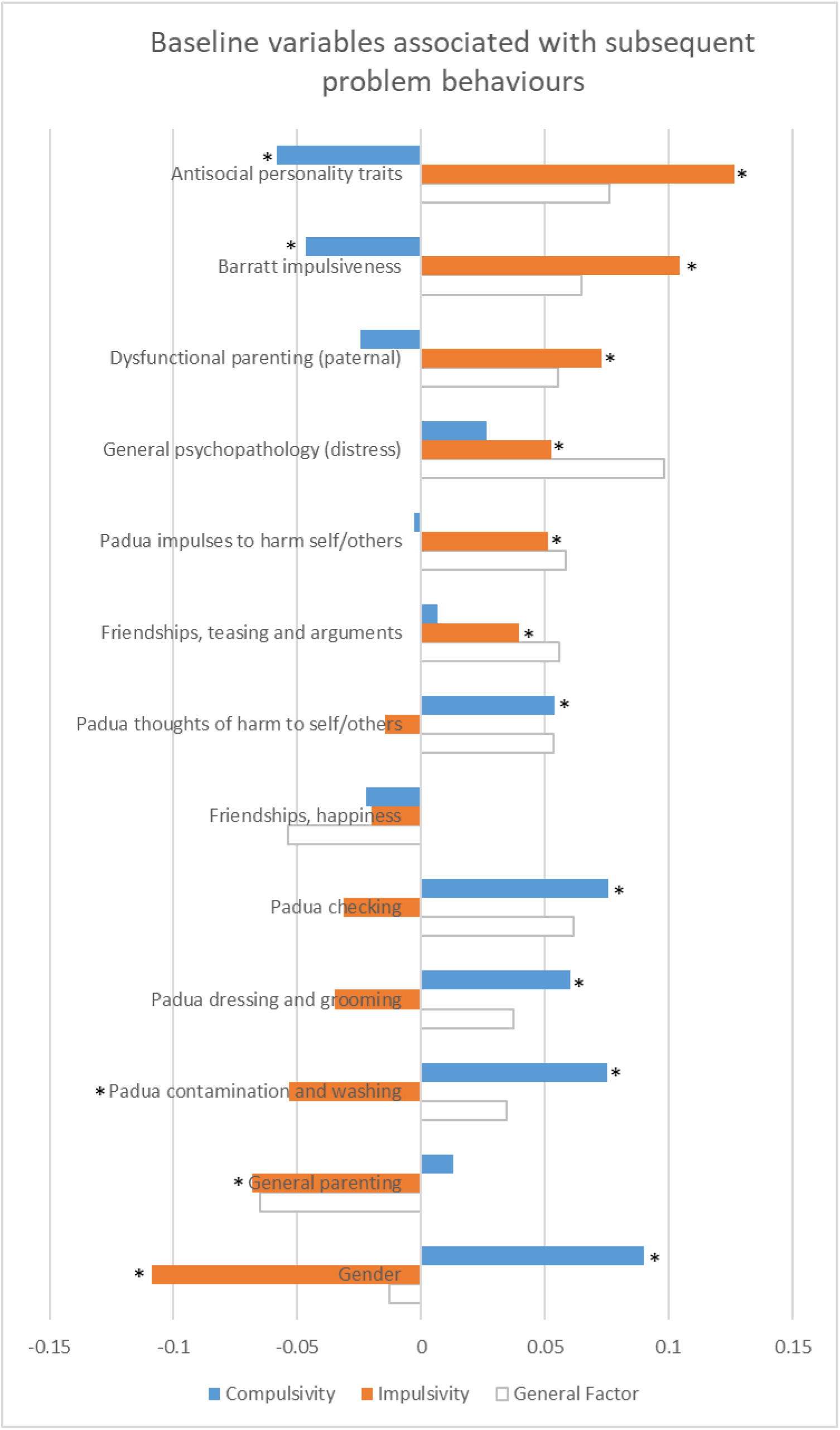
Standardized model coefficients for variables statistically explaining later problem behaviors (orange: impulsive problems; blue: compulsive problems). * indicates p<0.05 significant by rigorous statistical correction (bootstrap) for impulsive and compulsive problem behaviors. (For reference, model coefficients for the General Factor are shown in light grey outline; these were all significant by bootstrap except for gender [asterisks not shown]).

**Table 2.** Summary of Fit Statistics for the Different Competing Confirmatory Factor Analysis Models of the Impulsive Compulsive Behaviour Checklist (ICBC) in the NSPN Cohort

| | Model | *df* | $\chi^2$ | *p* | RMSEA (90%CI) | CFI | WRMR |
| --- | --- | --- | --- | --- | --- | --- | --- |
| 1 | Correlated Two-Factor <sup>1</sup> | 484 | 872.282 | <.001 | .035 (.032 - .039) | .960 | 1.146 |
| 2 | Orthogonal Two-Factor <sup>1</sup> | 485 | 3996.889 | <.001 | .106 (.103 - .109) | .643 | 3.507 |
| 3 | One-Factor <sup>1</sup> | 485 | 1187.035 | <.001 | .047 (.044 - .051) | .929 | 1.403 |
| 4 | Bi-Factor <sup>1</sup> | 461 | 647.001 | <.001 | .025 (.020 - .029) | .981 | .886 |

*Note.* <sup>1</sup>Model included error covariances, all significant when corrected for multiple post hoc comparisons using the Benjamini-Hochberg False Discovery Rate (FDR) ( $p = .05$ ); Chi square test statistic ( $\chi^2 > .05$ exact fit), Root Mean Square Error of Approximation (RMSEA) ( $\epsilon < .05$ close approximate fit; $\epsilon = .05 - .08$ close approximate fit; $\epsilon = .08 - 1.0$ reasonable approximate fit); Comparative Fit Index (CFI) ( $> .90$ reasonable fit; $> .95$ good fit); Weighted Root Mean Residual (WRMR) ( $< .950$ good fit)

**Table 3.** Standardized Loading Estimates for individual ICBC Items on the Disinhibition, Impulsivity, and Compulsivity Factors. All loading estimates were significant at p < .001, except where indicated. **p < .01, *p < .05.

| ICBC Items | Disinhibition | Impulsivity | Compulsivity |
| --- | --- | --- | --- |
| 1. Washing | .677 |  |  |
| 2. Smoking | .141* | .585 |  |
| 3. Collect | .519 |  |  |
| 4. Money | .295 |  | .375 |
| 5. Ordering | .626 |  | .588 |
| 6. Shopping | .697 |  |  |
| 7. List | .586 |  | .496 |
| 8. Counting | .705 |  | .385 |
| 9. Grooming | .790 |  |  |
| 10. Routines | .674 |  | .366 |
| 11. Repeating | .684 |  | .234 |
| 12. Exercising | .629 |  |  |
| 13. Betting | .312 | .443 |  |
| 14. Hair Picking | .427 |  |  |
| 15. Lying | .602 | .405 |  |
| 16. Sexual | .592 | .498 |  |
| 17. Alcohol | .379 | .526 |  |
| 18. Planning | .608 |  | .451 |
| 19. Drug | .168* | .775 |  |
| 20. Cleaning | .543 |  | .422 |
| 21. Verbal | .625 | .489 |  |
| 22. Violence | .605 | .504 |  |
| 23. Swearing | .532 | .470 |  |
| 24. Checking Locks | .515 |  | .388 |
| 25. Checking Mirror | .751 |  |  |
| 26. Driving | .363 | .201** |  |
| 27. Medication | .655 | .214** |  |
| 28. Aggression | .637 | .429 |  |
| 29. Social | .616 |  |  |
| 30. Rules | .636 |  | .257 |
| 31. Injury | .474 | .310 |  |
| 32. Rewriting | .678 |  | .295 |
| 33. Tattooing | .595 | .437 |  |

### Baseline measures associated with subsequent trans-diagnostic problem behaviours

The optimal PLS model relating baseline variables to current impulsive-compulsive behaviours (ICBC latent scores) accounted for 37.4% of variation in the explanatory (X) measures and 11.6% of variation in later ICBC problem behaviour scores (Y). Variables that were important in the model (passing Variable Importance Parameter [VIP] threshold of 0.8) are shown in **Figure 1**. The following baseline variables were each significantly associated with higher subsequent impulsive problem behaviours: antisocial personality traits, impulsive traits (Barratt Impulsiveness Scale), dysfunctional perceived parenting scores (both general parenting and paternal parenting scores), general psychopathology/distress (K10), male gender, impulse to harm self/others (Padua Inventory subscore), and teasing/arguments in friendships. Impulsive problem behaviours were also significantly predicted by previous lower contamination obsessions and washing compulsions (Padua Inventory subscore). Baseline variables that were significantly associated with higher subsequent compulsive problem behaviours were: obsessional thoughts of harm to self/others (Padua Inventory subscore), compulsive checking (Padua subscore), dressing/grooming compulsions (Padua Inventory subscore), contamination obsessions and washing compulsions (Padua Inventory subscore), and female gender. Higher levels of compulsive problem behaviours were also significantly predicted by previous lower antisocial personality traits, and lower impulsive traits (Barratt Impulsiveness Scale). The following variables were statistically unimportant in the model and were excluded (VIP < 0.8): age at entry, ethnic group, family education level, current or past mental health or behavioural problem (parent/guardian report), history of head trauma (parent/guardian report), history of perinatal complications (parent/guardian report), and history of any medical conditions (parent/guardian report).

[FIG. 1 AROUND HERE PLEASE]

### Current measures associated with trans-diagnostic problem behaviours

The optimal PLS model relating current variables to current impulsive-compulsive behaviours (ICBC latent scores) accounted for 48.0% of variation in the explanatory measures (X) and 32.7% of variation in ICBC problem behaviour scores (Y). Variables that were important in the model (passing Variable Importance Parameter [VIP] threshold of 0.8) are shown in **Figure 2**. The following measures were each significantly associated with higher impulsive problem behaviours: antisocial personality traits, impulsive traits (Barratt Impulsiveness Scale), impulses to harm self/others (Padua Inventory subscore), and male gender. Higher impulsive problem behaviours were also significantly associated with fewer OCPD traits, lower dressing/grooming compulsions (Padua Inventory subscore), and lower contamination obsessions and washing compulsions (Padua Inventory subscore). The following measures were each significantly associated with higher compulsive problem behaviours: checking compulsions (Padua Inventory subscore), thoughts of harm to self/others (Padua Inventory subscore), dressing/grooming compulsions (Padua Inventory subscore), contamination obsessions and washing compulsions (Padua Inventory subscore), threat perception (OBQ), compulsive traits (CHIT), perfection and intolerance of uncertainty (OBQ), importance and control of thoughts (OBQ), OCPD traits, and female gender. Lower impulses to harm self/others (Padua Inventory subscore) was significantly associated with higher compulsive problem behaviours. The following variables were statistically unimportant in the model and were excluded (VIP < 0.8): age, ethnic group, education levels, and general psychopathology (distress).

**Figure 2.**
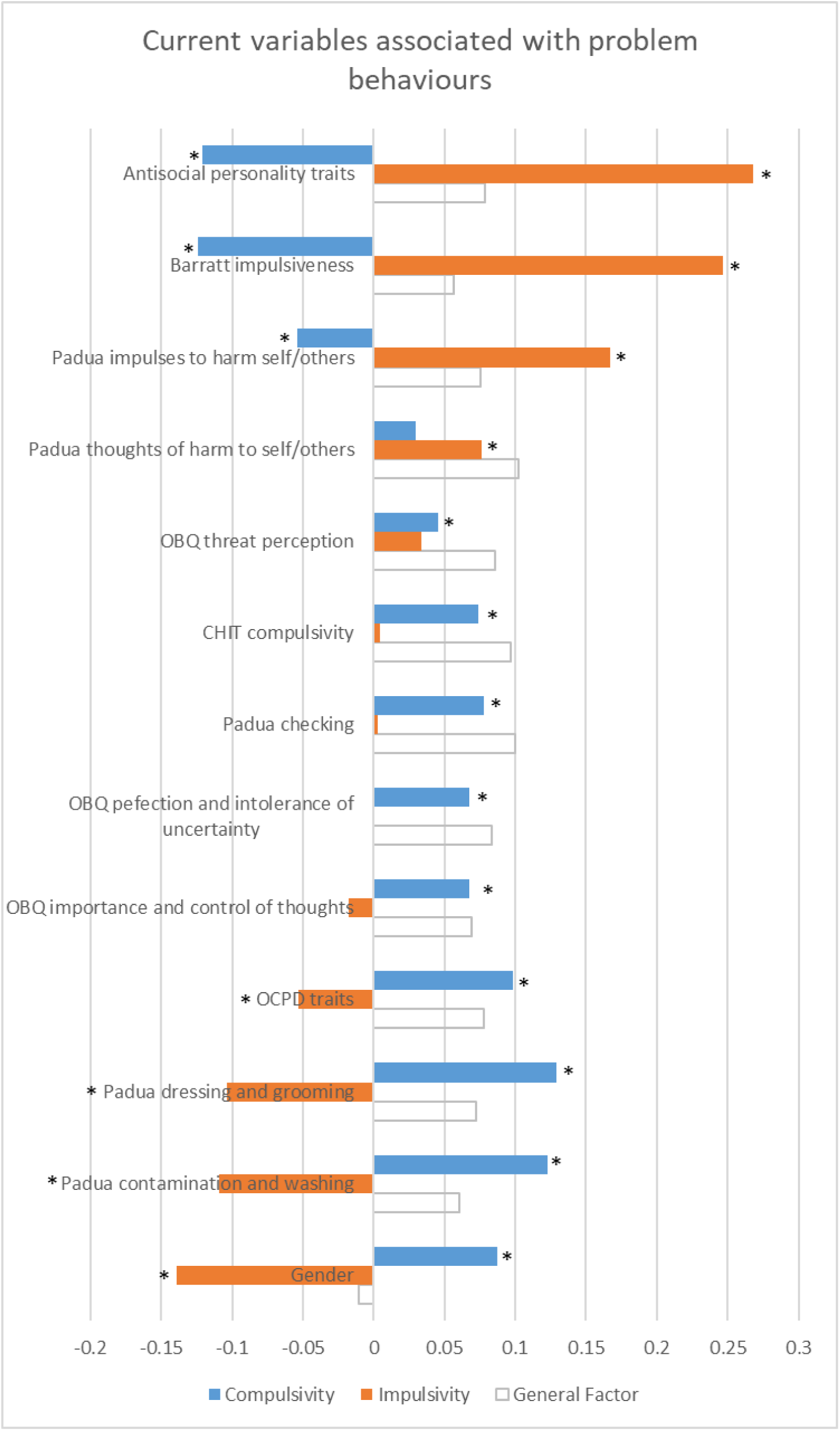
Standardized model coefficients for variables statistically explaining current problem behaviors (orange: impulsive problems; blue: compulsive problems). * indicates p<0.05 significant by rigorous statistical correction (bootstrap) for impulsive and compulsive problem behaviors. (For reference, model coefficients for the General Factor are shown in light grey outline; these were all significant by bootstrap except for gender [asterisks not shown]).

A conceptual schematic of the relationships between variables of interest and latent phenotype scores is provided in **Figure 3**.

**Figure 3.**
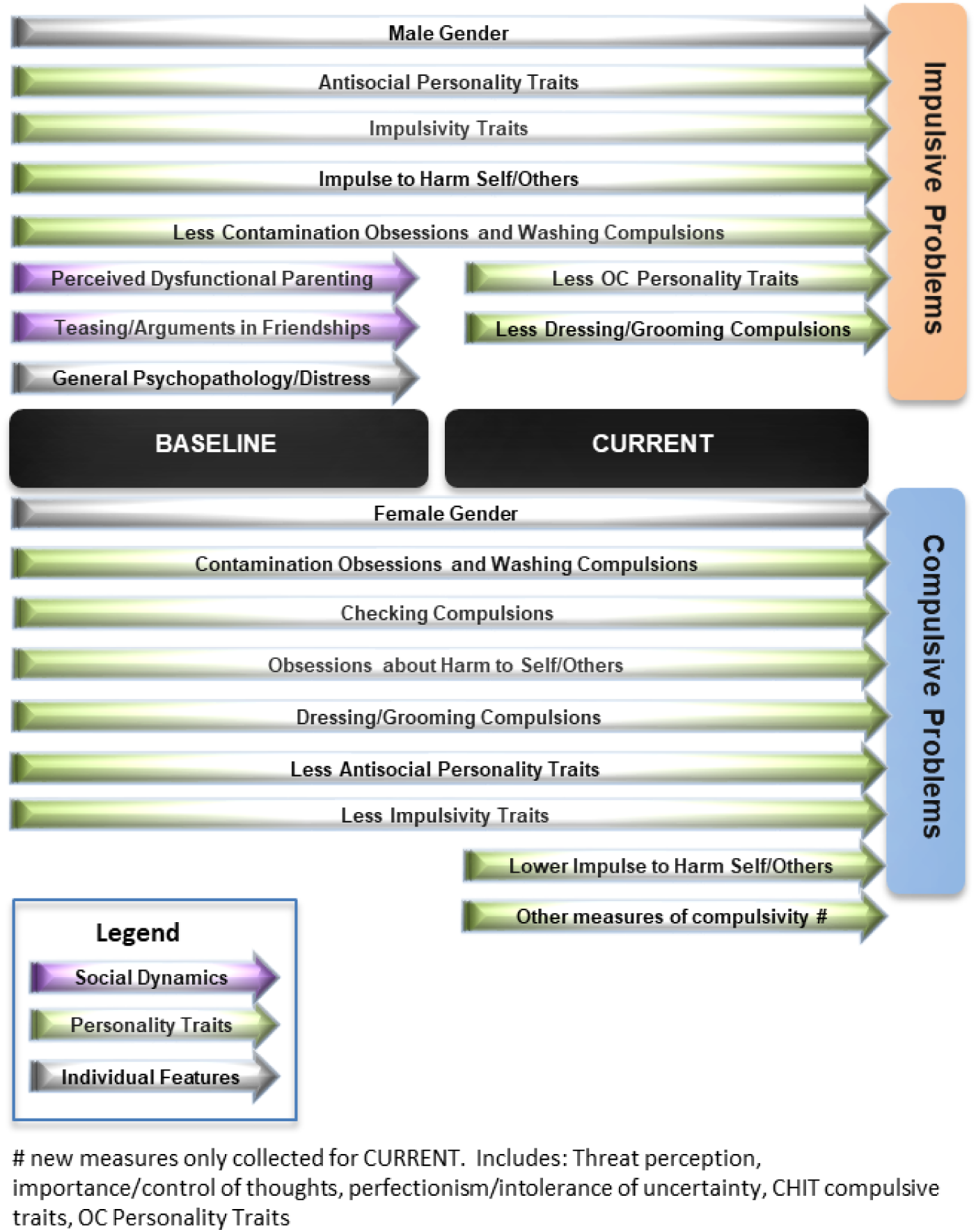
Variables significantly (p<0.05 bootstrap in PLS models) mapping onto trans-diagnostic phenotypes of impulsive and compulsive problem behaviors. OC = obsessive-compulsive.

[FIG. 2 AROUND HERE PLEASE]

[FIG. 3 AROUND HERE PLEASE]

## Discussion

This study examined latent phenotypes of impulsive and compulsive problem behaviours in young adults. We demonstrated that these behaviours were best conceptualised within a bi-factor model, which included two underlying latent phenotypes corresponding to impulsive and compulsive problems respectively (Guo et al., 2017) as well as a general disinhibition factor. By modelling this additional general factor, we could quantify trans-diagnostic phenotypes that were separable and not due (for example) to common measurement variance (Reise, 2012a). Predictors of the later emergence of these phenotypes were identified (**Figure 3**), incorporating relevant measures of personality, parenting, and friendship experiences. The statistical approach used (Partial Least Squares) allowed intrinsic control for co-relationships between the important explanatory variables of interest. We did not identify any significant effects of age or education levels on occurrence of problem impulsive or compulsive behaviours.

Personality-related questionnaires provide rich dimensional measures, and are typically developed for use in normative settings, as well as in patient populations (Stanford et al., 2016; Sanavio, 1988). Personality measures here showed the clearest differentiation in statistically predicting later trans-diagnostic occurrence of impulsive and compulsive problem behaviours (**Figure 3**). Higher baseline scores on the Barratt Impulsivity Scale (Stanford et al., 2016), and the Antisocial Process Screening Device (Vitacco et al., 2003), each significantly predicted higher subsequent impulsivity scores derived from the ICBC (an independent instrument); and contrariwise, each of these two baseline personality scores significantly predicted lower subsequent compulsive problem behaviours.

As anticipated, most of the Padua obsessive-compulsive Inventory subscores (Sanavio, 1988) were associated with later emergence of compulsive but not impulsive problem behaviours, especially so for archetypal contamination obsessions and washing compulsions (for further discussion of Padua Inventory results see **Supplement**). We view scores on the Padua Inventory as personality traits in that they are measurable in the general population along a continuum. In fact the instrument was originally developed for community use (Sanavio, 1988); this is in contrast to clinical measures of OCD, such as the Yale-Brown Obsessive Compulsive Scale (Y-BOCS), which are unsuitable for phenotyping research due to a high likelihood of zero score in those without formal OCD (Goodman et al., 1989).

While the above associations are largely expected, in view of the conceptual domains they encompass, the results also indicate important, differential contribution of earlier personality traits known to be partly heritable (Niv et al., 2012; Tuvblad et al., 2014; Jonnal et al., 2000) to the ultimate manifestation of an extensive array of 33 adult problem behaviours. This pattern of association for impulsive, antisocial, and OC traits was also replicated cross-sectionally, but extending to a broader range of compulsivity instruments not originally included at baseline in the NSPN cohort (**Figure 3**).

It is well established that among environmental factors relevant to later risk of mental health problems, parenting and friendship networks are particularly important in young people, along with general distress. For example, family and friendship support during early adolescence negatively predicted risk of later adolescent depressive symptoms arising as a function of childhood adversity (van Harmelen et al., 2016). Here we show that worse perceived general parenting, higher levels of perceived dysfunctional parenting (reported by the study participant in relation to their father), and higher levels of general distress (K10 instrument) were specifically associated with later emergence of impulsive but not compulsive problems (**Figure 3**). Furthermore, participants’ earlier experiences of teasing and arguments within their friendship network, and not other aspects of friendship (as measured), were also significantly associated with later impulsive but not compulsive problem behaviours. Family functioning is an important determinant of specific types of impulsive problems (such as non-suicidal self-injury) (Cassels et al., 2018). Positive relationships between dysfunctional parenting (especially harshness and psychological control) and externalising symptoms have been shown in prior meta-analysis, in young people (Pinquart, 2017). In a longitudinal population-based study of adopted and non-adopted adolescents, parent-child conflict was associated with subsequent impulsive (‘acting-out’) behaviours (Klahr et al., 2011a; Klahr et al., 2011b).

We found a dissociation in the influence of gender on impulsive and compulsive trans-diagnostic phenotypes: male gender was significantly associated with impulsive problem behaviours (viewed trans-diagnostically) whereas female gender was significantly associated with compulsive problem behaviours. Impulsive problems (such as aggression and criminality) are more common in males in much of the literature (Chamorro et al., 2012), findings which may be due to males having higher sensation seeking and risk-taking (Cross et al., 2011). Several previous cross-sectional studies found that female gender is associated with higher obsessive-compulsive traits on the Padua Inventory (Chamberlain et al., 2016; Sanavio, 1988). These results hint at sexually divergent processes in the development of impulsive versus compulsive problems, which are not due (for example) to confounding gender differences in personality traits, parenting, or friendships, which were controlled for in the statistical modelling. Our data also accord well with findings from literature in other areas, particularly internalizing-externalizing studies in young people into adulthood, which commonly report a similar gender discrepancy (Wilhelm, 2014).

The bi-factor modelling approach is increasingly adopted in psychiatry research (Castellanos-Ryan et al., 2016; Hankin et al., 2016; Caspi et al., 2014; Patalay et al., 2015; Stochl et al., 2015). Here, we demonstrate that a bi-factor model yielded superior fit across all metrics for the fractionation of trans-diagnostic phenotypes of impulsive and compulsive problem behaviours. To measure the phenotypes of interest, we used responses from the ICBC, which examined a broad range of 33 behaviours in a single, convenient questionnaire. The nature of the general latent factor, within the bi-factor model, itself is open to interpretation, as in other studies. The general factor was associated with all study measures excepting gender, suggesting that it may relate to general disinhibition (general tendencies towards failure to suppress behaviours). We believe it unlikely that this general factor reflects only common response bias because its relationship with individual ICBC items was variable. It is intended to examine biological substrates of this general disinhibition factor in future work.

There are several limitations in relation to this study. Rigorous demonstration of causality has multiple scientific requirements and cannot be shown definitively within a study design as deployed here. Nonetheless, impulsive, dissocial, and compulsive traits bore strong relationships with their respective expected problem behaviours (measured using a separate questionnaire) both longitudinally and cross-sectionally, showing a sustained and meaningful association. The original recruitment methodology for NSPN was designed to be epidemiologically representative (i.e. stratified based on census statistics) (Kiddle et al., 2017). However, the current study participants provided data in 2018 and thus may not be fully representative of the original NSPN cohort. Our current sample had mean impulsive and compulsive scores (Barratt Impulsivity Scale and Padua Inventory) similar to those reported in previous normative studies (Burns et al., 1996; Steinberg et al., 2013). We measured relatively ‘high level’ trans-diagnostic phenotypes closely allied to overt problems, but the phenotypic approach should ultimately be applied to a range of measures incorporating also cognitive and biological (genetic, imaging) parameters. Lastly, the best fit statistical model for data is contingent on the particular nature and range of measures used; because a given model offers a superior fit, it does not follow that other models have no utility.

In summary, impulsive and compulsive problem behaviours were fractionated trans-diagnostically using a bi-factor model, which yielded the best fit versus other alternative models suggested by the extant literature. Significant baseline predictors of later problem impulsive and compulsive trans-diagnostic phenotypes were identified, using a powerful statistical technique well-suited to large datasets where items are likely to be correlated. Personality-traits were important determinants of impulsive and compulsive problem behaviours, whereas parenting and friendship experiences significantly predicted impulsive problems specifically, and gender specific effects were found (male gender, higher impulsive problems; female gender, higher compulsive problems). This study highlights the potential utility of the trans-diagnostic approach in psychiatry, as applied to impulsivity and compulsivity. The heritability and biological substrates of latent impulsivity, compulsivity, and general model factors merits exploration. For example, if the trans-diagnostic phenotype of compulsivity has higher heritability than impulsivity, this may account for the relatively lower impact of the earlier social milieu on the former. The finding that aspects of parenting and friendships differentially relate to impulsive as opposed to compulsive problems may have implications for tailoring early interventions and targeting vulnerable individuals to avert the development of later problematic behaviours. In particular, such trans-diagnostic approaches can be used to identify candidate risk factors for a broad range of psychiatric symptoms. These risk factors (such as elements of personality, friendship support, and parenting) could in future be targeted for modification in people at risk of impulsive and compulsive problems, in order to avert the development of these impairing pathologies. This would potentially shift the emphasis in psychiatry more towards prevention (Furber et al., 2017), as well as treatment; a shift that has been deployed with success in other areas of healthcare, and in other areas of psychiatry outside the impulsive-compulsive sphere.

## Potential conflicts of interest

Dr. Chamberlain consults for Cambridge Cognition and Shire. Dr. Grant has received research grants from NIDA, National Center for Responsible Gaming, American Foundation for Suicide Prevention, and Forest and Roche Pharmaceuticals Dr. Grant receives yearly compensation from Springer Publishing for acting as Editor-in-Chief of the Journal of Gambling Studies and has received royalties from Oxford University Press, American Psychiatric Publishing, Inc., Norton Press, Johns Hopkins University Press, and McGraw Hill. Jeggan Tiego was supported by National Health and Medical Research Council (NHMRC) project grants 1050504 and 1146292. Dr. Fontenelle is supported by the “Fundação Carlos Chagas Filho de Amparo à Pesquisa do Estado do Rio de Janeiro (FAPERJ) under E-26/010.001411/2015; and “Conselho Nacional de Desenvolvimento Científico e Tecnológico” (CNPq)] under grant 308237/2014-5. Dr. Goodyer consults for Lundbeck; is supported by a Wellcome Trust Strategic Award; and is Chairperson of and scientific advisor to the Peter Cundill Centre for Youth Depression Research, Centre for Addictions and Mental Health, University of Toronto. Dr. Yücel was supported by a National Health and Medical Research Council of Australia Fellowship (#APP1117188) and the David Winston Turner Endowment Fund. Dr Rebecca Segrave was supported by the David Winston Turner Endowment Fund. The other authors report no conflicts of interest or disclosures. Dr Hauser is supported by the Jacobs Foundation. Dr Dolan is supported by a Wellcome Investigator Award and by the Max Planck Society. Dr. Bullmore is employed half-time by GSK and holds stock in GSK.

### Acknowledgement

The authors would like to thank the NSPN study team, particularly Gita Prabhu and Laura Villis, for invaluable assistance in setting up the study; and would like to thank the study participants. This research was supported by the NIHR Cambridge Biomedical Research Centre (BRC).

## Clinical trials registration

not applicable.

## Previous presentations

none.

