## Supplement for "Fractionation of impulsive and compulsive trans-diagnostic phenotypes and their longitudinal associations"

**Supplementary Online File – Chamberlain et al.**

**Supplementary Table 1. Number and percentage of sample endorsing each problematic behaviour, ranked by highest to lowest. For original item descriptions and validation see (Guo et al., 2017).**

|  | Problematic (often/always) | |
| --- | --- | --- |
| ICBC Scale item (behaviour) | N | % |
| impcomp_7_list | 97 | 14.8 |
| impcomp_5_ordering | 88 | 13.5 |
| impcomp_4_money | 85 | 13.0 |
| impcomp_6_shopping | 81 | 12.4 |
| impcomp_18_planning | 73 | 11.2 |
| impcomp_23_swearing | 73 | 11.2 |
| impcomp_29_social_networking | 72 | 11.0 |
| impcomp_24_checkinglocks | 68 | 10.4 |
| impcomp_8_counting | 61 | 9.3 |
| impcomp_12_exercising | 57 | 8.7 |
| impcomp_3_collect | 56 | 8.6 |
| impcomp_25_checkingmirror | 56 | 8.6 |
| impcomp_32_rewriting | 53 | 8.1 |
| impcomp_10_routines | 51 | 7.8 |
| impcomp_9_grooming | 45 | 6.9 |
| impcomp_1_washing | 41 | 6.3 |
| impcomp_14_hairpicking | 39 | 6.0 |
| impcomp_11_repeating | 37 | 5.7 |
| impcomp_17_alcohol | 35 | 5.4 |
| impcomp_30_rules | 34 | 5.2 |
| impcomp_20_cleaning | 32 | 4.9 |
| impcomp_16_sexual | 31 | 4.7 |
| impcomp_15_lying | 26 | 4.0 |
| impcomp_21_verbal | 24 | 3.7 |
| impcomp_31_injury | 18 | 2.8 |
| impcomp_2_smoking | 17 | 2.6 |
| impcomp_26_driving | 15 | 2.3 |
| impcomp_19_drug | 12 | 1.8 |
| impcomp_33_tattooing | 12 | 1.8 |
| impcomp_27_medication | 11 | 1.7 |
| impcomp_22_violence | 10 | 1.5 |
| impcomp_28_aggression | 10 | 1.5 |
| impcomp_13_betting | 8 | 1.2 |

**Supplementary Table 2. Factor analysis for Family measures; baseline FAD, PPQ, and MOPs. Varimax rotation. Factor analysis yielded an optimal three-factor solution corresponding broadly to general parenting, paternal parenting, and maternal parenting.**

|  | **Factor 1** | **Factor 2** | **Factor 3** |
| --- | --- | --- | --- |
| fad_01_planning_activities_B1 | -0.547238 | 0.208334 | 0.076752 |
| fad_02_cant_talk_sadness_B1 | -0.672641 | 0.129103 | 0.058860 |
| fad_03_feel_accepted_B1 | 0.472165 | -0.264937 | -0.080512 |
| fad_04_dont_get_along_B1 | -0.565752 | 0.245882 | 0.180767 |
| fad_05_express_feelings_B1 | 0.674635 | -0.133244 | -0.066200 |
| fad_06_crisis_support_B1 | 0.601494 | -0.152014 | -0.031224 |
| fad_07_avoid_fears_B1 | -0.526606 | 0.081896 | 0.043102 |
| fad_08_solve_problems_B1 | 0.558286 | -0.207280 | -0.153954 |
| fad_09_confide_each_other_B1 | 0.662436 | -0.093072 | -0.026798 |
| fad_10_bad_feelings_B1 | -0.471003 | 0.298365 | 0.131157 |
| fad_11_accepted_B1 | 0.507883 | -0.241655 | -0.111908 |
| fad_12_decisions_problem_B1 | -0.487254 | 0.248221 | 0.099861 |
| apq_01_good_job_B1 | 0.609592 | -0.149168 | -0.228817 |
| apq_02_threaten_punish_B1 | -0.265094 | 0.180640 | 0.111741 |
| apq_03_fails_note_B1 | -0.288870 | 0.102757 | 0.081039 |
| apq_04_games_B1 | 0.625299 | -0.150003 | -0.195314 |
| apq_05_talk_out_punish_B1 | -0.080303 | 0.068991 | -0.020814 |
| apq_06_ask_school_B1 | 0.516450 | -0.114427 | -0.270524 |
| apq_07_stays_out_B1 | -0.144916 | 0.097129 | 0.058304 |
| apq_08_homework_B1 | 0.476666 | -0.112805 | -0.128946 |
| apq_09_compliment_B1 | 0.646527 | -0.145398 | -0.287235 |
| apq_10_praise_B1 | 0.549897 | -0.143812 | -0.184160 |
| apq_11_out_friends_B1 | -0.402493 | 0.087886 | 0.107516 |
| apq_12_punish_early_B1 | -0.045177 | 0.070352 | 0.012837 |
| apq_13_spank_B1 | -0.143259 | 0.230561 | 0.241285 |
| apq_14_slap_B1 | -0.235086 | 0.310520 | 0.276571 |
| apq_15_hit_B1 | -0.168838 | 0.220696 | 0.208459 |
| mops_mum01_overprotective_B1 | -0.117161 | 0.121324 | -0.139883 |
| mops_mum02_abusive_B1 | -0.279530 | 0.165638 | 0.569752 |
| mops_mum03_overcontrolling_B1 | -0.235632 | 0.196838 | 0.218999 |
| mops_mum04_guilty_B1 | -0.250222 | 0.269687 | 0.456317 |
| mops_mum05_ignored_B1 | -0.169700 | 0.186528 | 0.677657 |
| mops_mum06_critical_B1 | -0.296096 | 0.218219 | 0.409755 |
| mops_mum07_unpredictable_B1 | -0.180801 | 0.158012 | 0.603880 |
| mops_mum08_uncaring_B1 | -0.146691 | 0.018113 | 0.829674 |
| mops_mum09_violent_B1 | -0.213518 | 0.177831 | 0.596283 |
| mops_mum10_rejecting_B1 | -0.105363 | 0.054073 | 0.773359 |
| mops_mum11_left_B1 | -0.232416 | 0.128328 | 0.515551 |
| mops_mum12_forget_B1 | -0.113307 | 0.004932 | 0.707009 |
| mops_mum13_uninterested_B1 | -0.150325 | -0.035921 | 0.722987 |
| mops_mum14_danger_B1 | -0.071321 | 0.137109 | 0.607757 |
| mops_mum15_unsafe_B1 | -0.101437 | 0.066390 | 0.649114 |
| mops_dad01_overprotective_B1 | -0.118675 | -0.095268 | -0.023414 |
| mops_dad02_abusive_B1 | -0.302794 | 0.615893 | 0.068391 |
| mops_dad03_overcontrolling_B1 | -0.296815 | 0.281799 | 0.090779 |
| mops_dad04_guilty_B1 | -0.291750 | 0.556320 | 0.129004 |
| mops_dad05_ignored_B1 | -0.193290 | 0.745416 | 0.157472 |
| mops_dad06_critical_B1 | -0.249017 | 0.549219 | 0.077189 |
| mops_dad07_unpredictable_B1 | -0.252191 | 0.666023 | 0.108111 |
| mops_dad08_uncaring_B1 | -0.150943 | 0.810779 | 0.157088 |
| mops_dad09_violent_B1 | -0.228340 | 0.563792 | 0.040966 |
| mops_dad10_rejecting_B1 | -0.123108 | 0.796905 | 0.197000 |
| mops_dad11_left_B1 | -0.199347 | 0.630235 | 0.172926 |
| mops_dad12_forget_B1 | -0.113729 | 0.744063 | 0.182330 |
| mops_dad13_uninterested_B1 | -0.168528 | 0.738255 | 0.148740 |
| mops_dad14_danger_B1 | -0.117250 | 0.759869 | 0.080733 |
| mops_dad15_unsafe_B1 | -0.154001 | 0.748807 | 0.086786 |
| ppq_01_quality_time_B1 | 0.659200 | -0.111140 | -0.133581 |
| ppq_02_attend_school_B1 | 0.553836 | -0.205185 | -0.264559 |
| ppq_03_affection_B1 | 0.647323 | -0.151684 | -0.187810 |
| ppq_04_get_me_B1 | 0.490038 | -0.241933 | -0.234575 |
| ppq_05_comforted_B1 | 0.741944 | -0.218714 | -0.203940 |
| ppq_06_angry_listen_B1 | 0.739782 | -0.139884 | -0.176330 |
| ppq_07_praised_B1 | 0.676847 | -0.185377 | -0.305175 |
| ppq_08_ideas_encouraged_B1 | 0.702899 | -0.208536 | -0.279036 |
| ppq_09_priority_B1 | 0.689143 | -0.201396 | -0.306553 |
| ppq_10_loved_B1 | 0.656038 | -0.259801 | -0.354993 |
| ppq_11_listened_to_B1 | 0.772309 | -0.228558 | -0.249767 |
| ppq_12_contact_B1 | 0.546596 | -0.206974 | -0.300136 |
| ppq_13_home_safe_B1 | 0.394675 | -0.388621 | -0.413940 |
| ppq_14_opinions_valued_B1 | 0.755413 | -0.200207 | -0.242450 |
| ppq_15_talked_important_B1 | 0.790570 | -0.128205 | -0.215853 |
| ppq_16_privacy_respected_B1 | 0.569107 | -0.247863 | -0.173495 |
| ppq_17_friends_welcomed_B1 | 0.386843 | -0.231163 | -0.262560 |
| ppq_18_clothes_B1 | 0.301213 | -0.194910 | -0.383426 |
| ppq_19_pocket_money_B1 | 0.336293 | -0.104612 | -0.150965 |
| ppq_20_ask_B1 | 0.541811 | -0.219459 | -0.250330 |
| ppq_21_achieve_B1 | 0.405041 | -0.104150 | -0.433269 |
| ppq_22_cared_unwell_B1 | 0.387813 | -0.184888 | -0.375585 |
| ppq_23_skills_B1 | 0.608445 | -0.208298 | -0.298424 |
| ppq_24_advice_B1 | 0.693763 | -0.204428 | -0.261923 |
| ppq_25_learn_school_B1 | 0.349574 | -0.132913 | -0.427536 |
| ppq_26_progress_B1 | 0.439963 | -0.147804 | -0.396667 |

**Supplementary Table 3. Factor analysis for baseline friendship questionnaire. Varimax rotation. Factor analysis yielded an optimal three-factor solution corresponding broadly to number of friend and happiness; confiding and understanding; and teasing/arguments.**

|  | **Factor 1** | **Factor 2** | **Factor 3** |
| --- | --- | --- | --- |
| fq_01_number_friends_B1 | 0.234507 | 0.741151 | -0.111058 |
| fq_02_see_friends_B1 | 0.363220 | 0.388374 | 0.133014 |
| fq_03_friends_understand_B1 | 0.809284 | 0.318706 | -0.220105 |
| fq_04_confide_in_friends_B1 | 0.665423 | 0.319904 | -0.152303 |
| fq_05_friends_tease_B1 | -0.057429 | -0.026811 | 0.666492 |
| fq_06_not_friends_tease_B1 | -0.113377 | -0.019231 | 0.524172 |
| fq_07_arguments_B1 | -0.039992 | -0.114816 | 0.670419 |
| fq_08_overall_happy_B1 | 0.414031 | 0.713689 | -0.183807 |

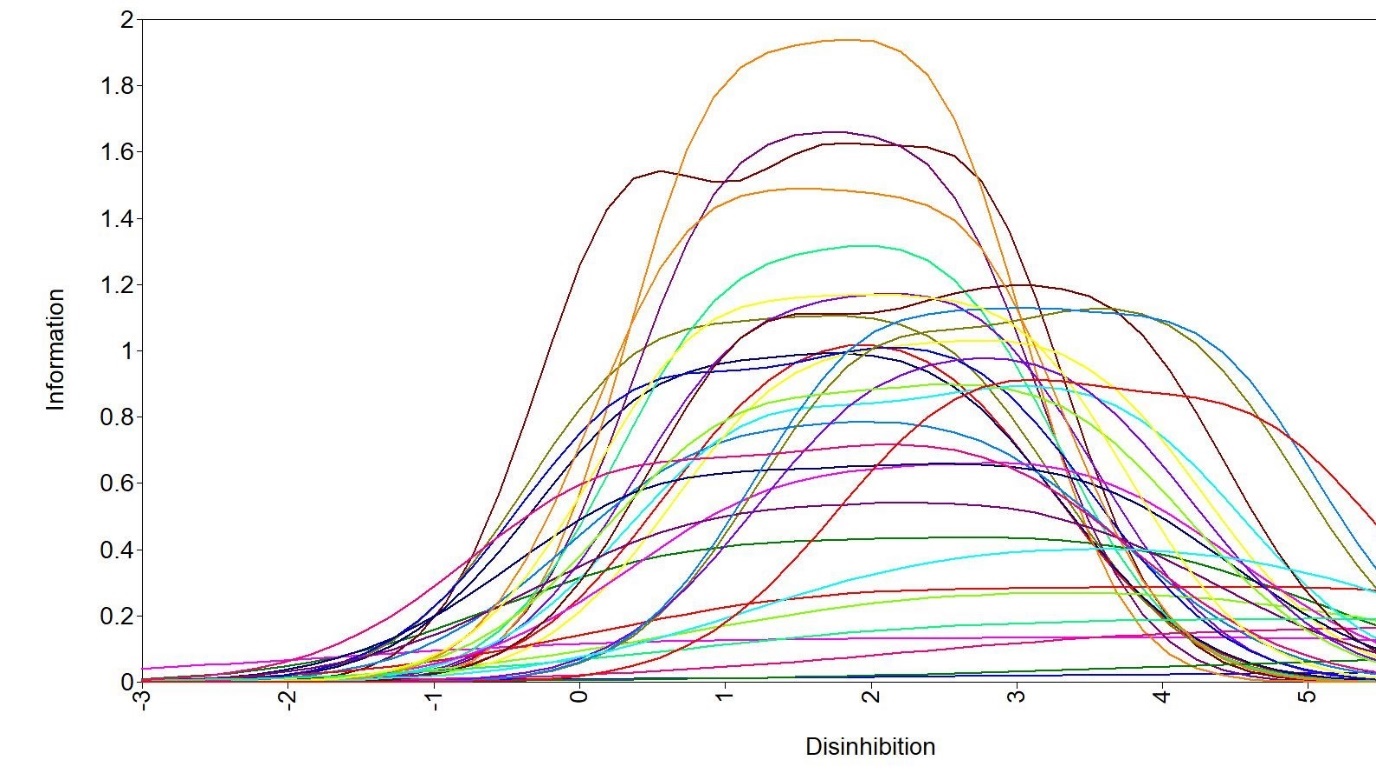

**Supplementary Figure 1. Item information functions for the 33 ICBC items in relation to the latent trait Disinhibition (θ). The information functions were distinct for different items, suggesting that the general Disinhibition factor from the bi-factor model of the ICBC was not attributable to response bias.**

**Additional Discussion regarding OC traits on the Padua Inventory**

As anticipated, most of the Padua obsessive-compulsive inventory subscores (Sanavio, 1988) were associated with later emergence of compulsive but not impulsive problem behaviours, especially so for archetypal contamination obsessions and washing compulsions. The one exception here was the ‘impulses to harm self/others’ Padua subscore, which was associated with impulsive problems, indicating that items originally designed to measure compulsivity in fact are endorsed by impulsive individuals, with implications for possible future revisions of this instrument. We view scores on the Padua inventory personality traits in that they are measurable in the general population along a continuum. In fact the instrument was originally developed for community use (Sanavio, 1988); this is in contrast to clinical measures of OCD, such as the Yale-Brown Obsessive Compulsive Scale, which are unsuitable for phenotyping research due to a high likelihood of zero score in those without formal OCD.
